# PyLossless: A non-destructive EEG processing pipeline

**DOI:** 10.1101/2024.01.12.575323

**Authors:** Scott Huberty, James Desjardins, Tyler Collins, Mayada Elsabbagh, Christian O’Reilly

## Abstract

EEG recordings are typically long and contain large amounts of data, making manual cleaning a time-consuming and error-prone task. Automated pre-processing pipelines can facilitate the efficient and objective extraction of artifacts, enabling standardized and reproducible analyses. However, automated pre-processing pipelines typically remove data considered artifact, and return a subset of irreversibly transformed signals. This approach obfuscates pre-processing decisions, and often makes it impossible to recover the original data or modify the pre-processing steps. Further, it complicates collaboration between research teams working on a common dataset, because different analyses may require specific pre-processing routines. Given the large amount of resources that are devoted to collecting EEG, tools that can help efficiently and transparently pre-process data are greatly needed. PyLossless addresses this need by creating a non-destructive, automated pre-processing pipeline that maintains the continuous EEG structure. It offers a user-friendly API, it is well documented, tested through continuous integration, easily deployable, and integrates with the popular MNE-Python environment. The pipeline further provides a browser-based quality control review (QCR) dashboard that allows researchers to visualize and edit the automated artifact annotations on sensors, time-periods, and independent components. The end product of PyLossless is a lossless annotated data state that can be shared and used with analysis-specific artifact rejection policies, allowing for an optimal balance between flexibility and standardization.

## Introduction

Electroencephalography (EEG) is a non-invasive technique widely used in neuroscience research and clinical applications. EEG recordings have a high temporal resolution, are typically long (up to multiple hours for sleep studies), and contain large amounts of data (sometimes more than 200 channels), making manual processing time-consuming and error-prone. Automated preprocessing pipelines can help to efficiently and objectively extract non-neural artifacts from EEG data, leading to more standardized and reproducible analyses. However, automated preprocessing pipelines typically remove data considered artifactual and return a subset of transformed signals [1–3]. This output is usually in the form of epochs, with sensors, time periods, or independent components containing artifacts automatically removed. This process can make it difficult for end-users to fully understand the decisions made regarding their data, i.e., which signals were kept or removed. Further, the output of such preprocessing pipelines is not well-suited for reusable and shareable datasets because different analyses might require alternative choices regarding epoch lengths, filtering properties, or independent component rejection. These issues are particularly relevant for EEG recordings from infants, children, and some clinical populations, which often contain a significant amount of stationary (e.g., EMG, ECG, blinks) and non-stationary (e.g., sweat, body movement) artifacts, often requiring expert review during preprocessing. As researchers invest significant resources in collecting EEG recordings, it is important to understand and correct, if necessary, the decisions made by an automated preprocessing pipeline. To this end, it is useful to split preprocessing into non-destructive and destructive phases, where the non-destructive phase adds annotations regarding artifactual signals and time periods to the data without altering the original signals. These annotations can be reviewed and amended before the destructive phase, using the saved annotations and a reproducible set of rules to generate preprocessed signals in a clear and non-ambiguous way. This separation allows for sharing an annotated dataset in a “ready-for-analysis” state without losing any data. These concepts are illustrated in Fig. 1.

**Fig 1.**
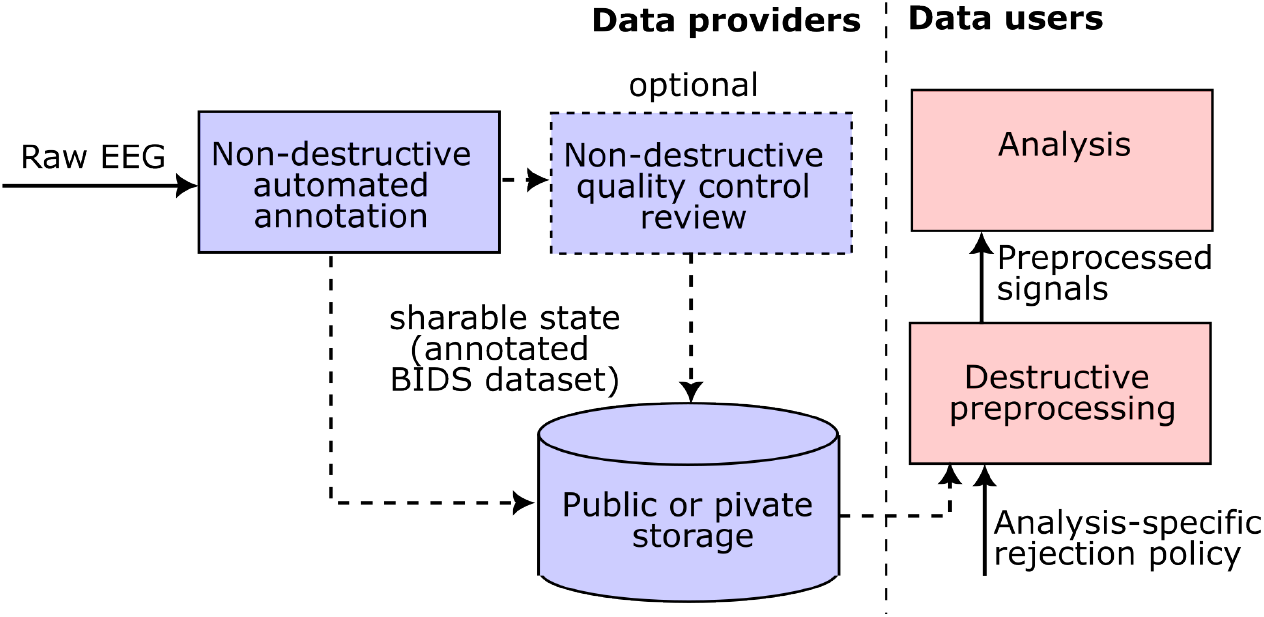
Proposed framework to support reproducible analyses with shared EEG datasets using a non-destructive EEG preprocessing process. This framework splits the preprocessing steps into non-destructive (purple) and destructive (red) phases. In the non-destructive phase, annotations are added to the continuous EEG signals, providing information about artifacts in the data. In the destructive phase, end-users can apply the annotations to the data in order to remove artifacts as needed for their respective analyses.

The EEG-IP-Lossless pipeline was previously proposed by Desjardins & colleagues (2021) [4] as a framework for the lossless preprocessing of EEG. This paper presented methods for annotating continuous EEG data for artifactual sensors, time periods, and independent components (ICs) in a non-destructive manner. This MATLAB pipeline accepted BIDS-compliant EEG datasets [5] as input and saved pipeline decisions as metadata (annotations) rather than irrevocably modifying the EEG signals. The approach proposed in this paper presents several advantages. First, this pipeline provides traceable annotations and mostly avoids destructive steps. This annotation-oriented design approach provides the users with complete flexibility with respect to how they wish to use these annotations for data preprocessing, making the pipeline output shareable and reusable for independent analyses. Second, this pipeline preprocessed data in its native, continuous form rather than in a derived epoched state. Keeping the annotation process at the level of continuous signals makes it more intuitive to review the decisions taken by the automated pipeline. Finally, this approach is much more appropriate for naturalistic paradigms, as recordings from such experiments are generally incompatible with epoching, given the ongoing and unstructured temporal relationship between naturalistic stimuli and responses. Unfortunately, although this paper pushed forward impactful concepts, its software implementation was overly complex and lacked robustness. Notably, its filtering and rereferencing processes produced some non-traceable transformation of original data, making it less than 100% lossless. Further, this MATLAB pipeline is no longer supported due to issues with software version incompatibilities and its complex deployment and usage.

PyLossless is a faithful implementation and extension of the MATLAB EEG-IP-Lossless pipeline, making it available to the large and growing scientific Python community. This pipeline, which we call PyLossless, significantly improves upon the MATLAB version by integrating closely with the MNE-Tools ecosystem, including MNE-Python for annotating continuous EEG data [6], MNE-BIDS for handling BIDS-compliant datasets [7], and MNE-ICALabel for automatically labeling independent components [8]. It also streamlines and clarifies the pipeline concepts, formalizes some new key ideas, ensures complete losslessness, and ensures software quality by adopting state-of-the-art DevOps (e.g., continuous integration, comprehensive test coverage, detailed documentation) and software engineering standards (e.g., interface segregation, componentization). Finally, PyLossless provides a quality control review (QCR) dashboard implemented with Plotly-Dash, which itself is built on top of React, making it easy to deploy with minimum dependencies. This agile deployment strategy for the QCR interface opens a wide range of applications, such as deployment on remote compute clusters for EEG processing in the context of Big Science projects. PyLossless is thoroughly tested via continuous integration to ensure that the pipeline methods continue to flag artifactual channels, time, and independent components as expected after future code updates. PyLossless does not seek backward compatibility or exact reproducibility with the MATLAB EEG-IP-Lossless pipeline but rather takes this opportunity to push its fundamental concepts forward significantly while validating that the output PyLossless is on par or superior with its defunct MATLAB counterpart.

## Materials and methods

### Licence and Dependencies

PyLossless is a Python 3 package distributed under the very permissive and open-science-friendly MIT license. Other than the libraries within Python’s core scientific stack (NumPy [9], SciPy [10], and Pandas [11]), PyLossless has minimal dependencies. Libraries in the MNE-Tools ecosystem (MNE-Python [6], MNE-BIDS [7], and MNE-ICLabel [8]) are used for handling EEG data, BIDS standardization, and labeling independent components, respectively. Given that MNE-Python has a large and growing user base and that it currently receives funding from the National Institutes of Health, the Chan Zuckerberg open-source initiative, and the European Research Council, it is likely that it will continue to serve as a strong foundation for the PyLossless pipeline. Plotly-Dash is a weak dependency required for users who wish to deploy the QCR dashboard but is not mandatory for using the automated functionality of the pipeline. As opposed to other popular Python GUI toolboxes like Qt which relies on C++ bindings and can sometimes present installation issues, Plotly-Dash calls JavaScript from Python to provide interactive graphics. PyLossless is welcoming of community contributions, and our documentation provides clear guidance on contributing, including the requirements for contributed code to be integrated into the pipeline.

### Documentation

Installation instructions, tutorials, implementation strategies, and the Application Programming Interface (API) are extensively documented [12]. The documentation is automatically built using Sphinx and deployed using GitHub actions, and tutorials are generated using Sphinx-Gallery. The documentation of the API adheres to the NumpyDoc style. PyLossless also hosts a GitHub discussion forum, where users can post questions about the pipeline.

### Testing and Continuous Integration

To prevent code regression, functional tests (smoke and unit tests) are systematically launched through GitHub continuous integration (CI) whenever a change is made to the PyLossless code base. CI also includes PEP8 compliance tests for code readability, and documentation building to ensure that PyLossless functions, classes, and methods are sufficiently detailed and that tutorials run without error.

### Design and Implementation

Below, we describe in detail each step of the PyLossless pipeline. Note that the operations in each step allow for some specific configuration by the user (for example, how many channels must be outliers for a time-period to be flagged as artifactual). For the sake of clarity, we will describe the operations using the pipeline’s current default parameters.

### Pipeline steps

PyLossless expects input data to be in BIDS format [2], but it also uses MNE-BIDS to provide helper functions for converting non-BIDS datasets to BIDS for preprocessing with PyLossless. BIDS compliance ensures that the pipeline can find and extract all necessary information. Before assessing data quality, PyLossless applies a robust average reference. Aside from helping to standardize the input data (i.e., different datasets may use various referencing schemes), this procedure controls for typical issues associated with single electrode reference, e.g., sensors that are further from the reference electrode will have a larger voltage variance. The robust average reference equalizes the voltage variance across sensors by subtracting the averaged signal across sensors from each channel. However, sensors contaminated with large artifacts are excluded from this average reference, so as not to inject artifacts into all electrodes. For a more detailed description regarding how the pipeline identifies electrodes to leave out of the robust average reference, see Supplementary Fig 9.

The different steps of the pipeline are shown in Fig 2. Note that these steps are said to “flag” sensors, ICs, or epochs/time periods. The concept of flagging refers to the creation of annotations that are saved as metadata at the end of the non-destructive phases of the pipeline (see Fig 1). These flags can then be used to tailor the destructive preprocessing phase, as defined for specific analyses by corresponding rejection policies specified as YAML files.

**Fig 2.**
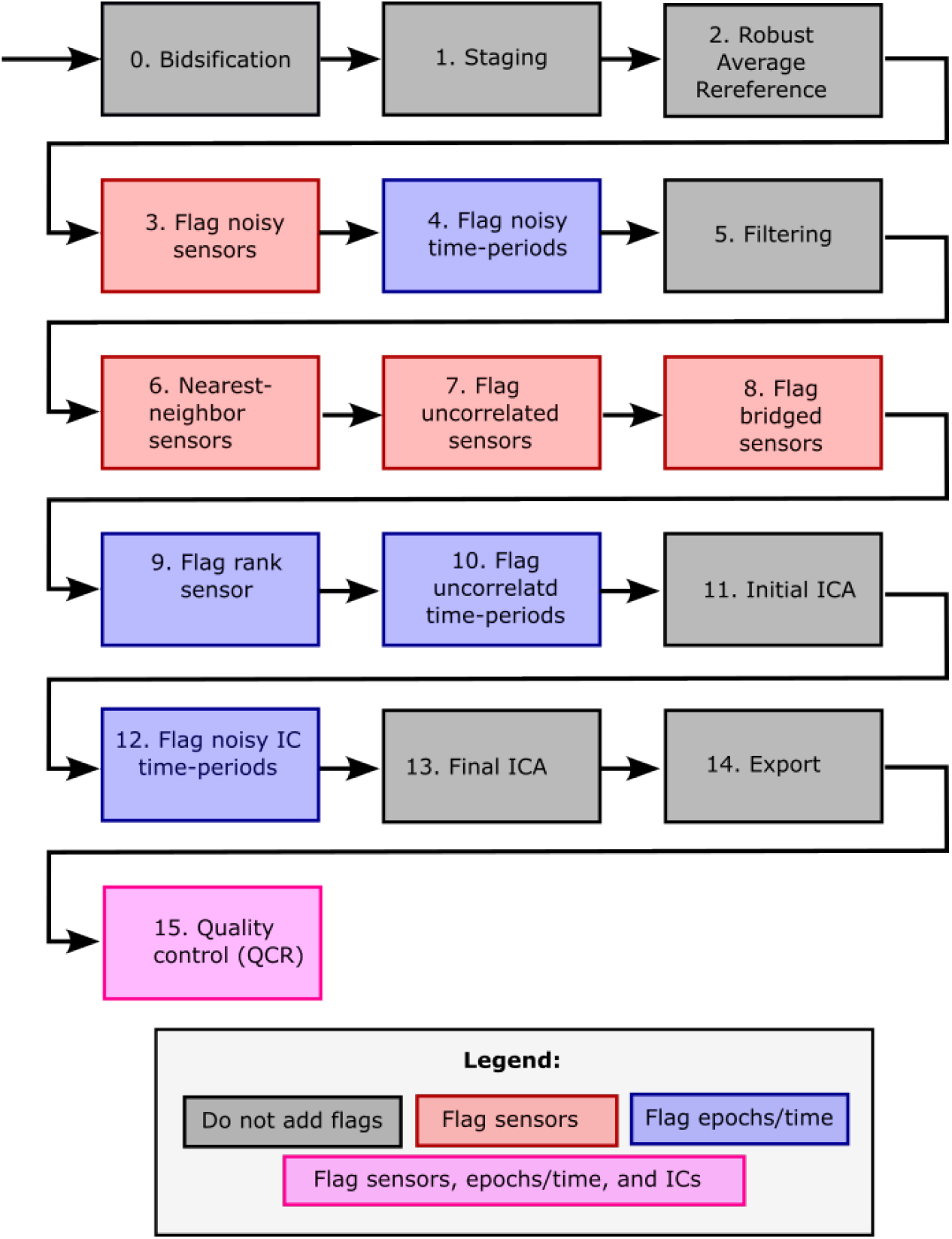
Steps of the PyLossless pipeline. The bidsification is numbered 0 because it is not part of the PyLossless pipeline itself, but rather a preliminary step required for non-BIDS-compliant datasets.

### Notation

To simplify the presentation, we will define some matrices and metrics that are used repeatedly throughout the pipeline. We denote a 3D Matrix of valid EEG data as

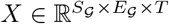 where *S*_*𝒢*_ and *E*_*𝒢*_ are the set of valid (i.e. not flagged) sensors and epochs, respectively, and T is the number of samples (i.e. time points). Note that this matrix is used throughout the pipeline and *S*_*𝒢*_ and *E*_*𝒢*_ are used to define its dimension to emphasize that as the pipeline proceeds and the set of valid sensors and epochs reduces, the size of this matrix changes. Let *s, e*, and *t* represent the sensors, epochs, and samples dimensions, respectively. We define the matrix containing the data for a single sensor *i* as 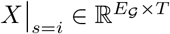, with *i* ∈ *S*_*𝒢*_. Likewise, a data associated with a single epoch *j* is noted 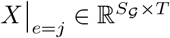, with *j* ∈ *E*_*𝒢*_.

The PyLossless functions flag sensors or epochs as artifacts based on rejection thresholds. Instead of applying a global rejection threshold to all sensors or epochs, most of the pipeline steps apply sensor-specific and epoch-specific thresholds. This is often preferable because, for example, different sensors may have differing baseline levels of voltage variance [3]. Sensor-specific thresholds for flagging epochs are defined as 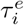. Here, subscript *i* identifies the sensor, whereas the superscript *e* denotes the fact that this threshold is applied to the distribution of data across epochs. Likewise, epoch-specific thresholds for flagging sensors are defined as 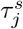.

The pipeline functions often calculate quantiles across the sensors or epochs. To simplify notation, we note quantiles as *Q*#^*dim*^ (i.e., *Q*75^*s*^ is the 75th quantile along the sensor dimension and the function *Q*75^*s*^(*X*) computes the 75th quantile along the sensors dimension *s* of matrix *X*, resulting in a matrix noted 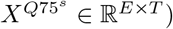.

Finally, throughout the text, we use capital letters for matrices and lower-case letters for scalars. For example, we can denote the data point for sensor *i*, epoch *j*, and time *k* as *X*|_*s*=*i*;*e*=*j*;*t*=*k*_ = *x*_*ijk*_ ∈ ℝ, and *X* = {*x*_*ijk*_}. We use superscripts to denote an operation across a dimension (i.e., aggregate), and we use subscripts to denote the index of a dimension (i.e., select). The pipeline performs series of such combinations of *select-and-aggregate* transformations of the original matrix *X* to identify various types of artifacts.

### Flag Noisy Sensors

Flag noisy sensors/time-periods steps use a metric based on the standard deviation to identify and flag sensors and time-periods with large, outlying voltage variance. In this step, we take the standard deviation of input matrix 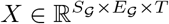 across the dimension samples, resulting in a 2D matrix 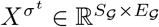. Then, we take the 50th and 75th quantile across the dimension sensor of 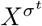. This operation results in two vectors of size *E*_*𝒢*_ :

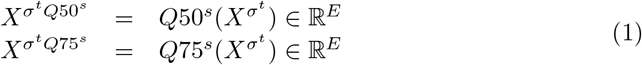

These vectors represent the 50th and 75th percentiles across sensors of the voltage dispersion across samples for each epoch. The difference between these two vectors (which we call the upper quantile range, UQR) is used to define a threshold for rejecting sensors:

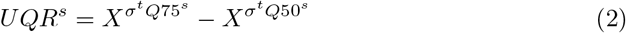

From here, the pipeline defines epoch-specific thresholds for flagging sensors. We multiply a constant *k* by *UQR*^*s*^ to define a measure for the spread of the right tail of the distribution of 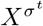 and add it to the median of 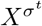 to obtain epoch-specific dispersion thresholds for detecting outlier sensors. Note that the constant *k* is a parameter that can be adjusted to control the sensitivity of the threshold. A larger value of *k* will result in a more conservative threshold:

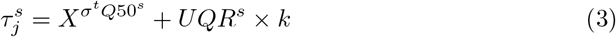

That is, 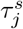 is the threshold used to detect outlier sensors specifically for epoch *j*.

Next, we compare our 2D dispersion matrix to the threshold vector 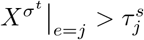 resulting in the indicator matrix 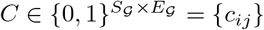 with

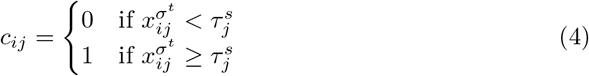

Each element of this matrix indicates whether a sensor *i* is an outlier at epoch *j*. To determine whether a sensor should be flagged for rejection, we average across the epochs dimension of our indicator matrix *C* and obtain:

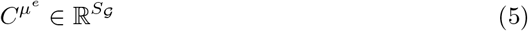

which is a vector of fractional numbers 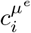 representing the percentage of epochs for which sensor *i* is an outlier. Next, we define a consensus threshold *τ*^*p*^ (*p* for percentile; default value: .20) as a cutoff point for determining if a sensor should be flagged. The sensor *i* is flagged as artifact if 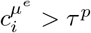. In other words, if a sensor is noisy in more than *τ*^*p*^ percent of epochs then it is flagged for rejection. These operations are represented schematically in Fig 3.

**Fig 3.**
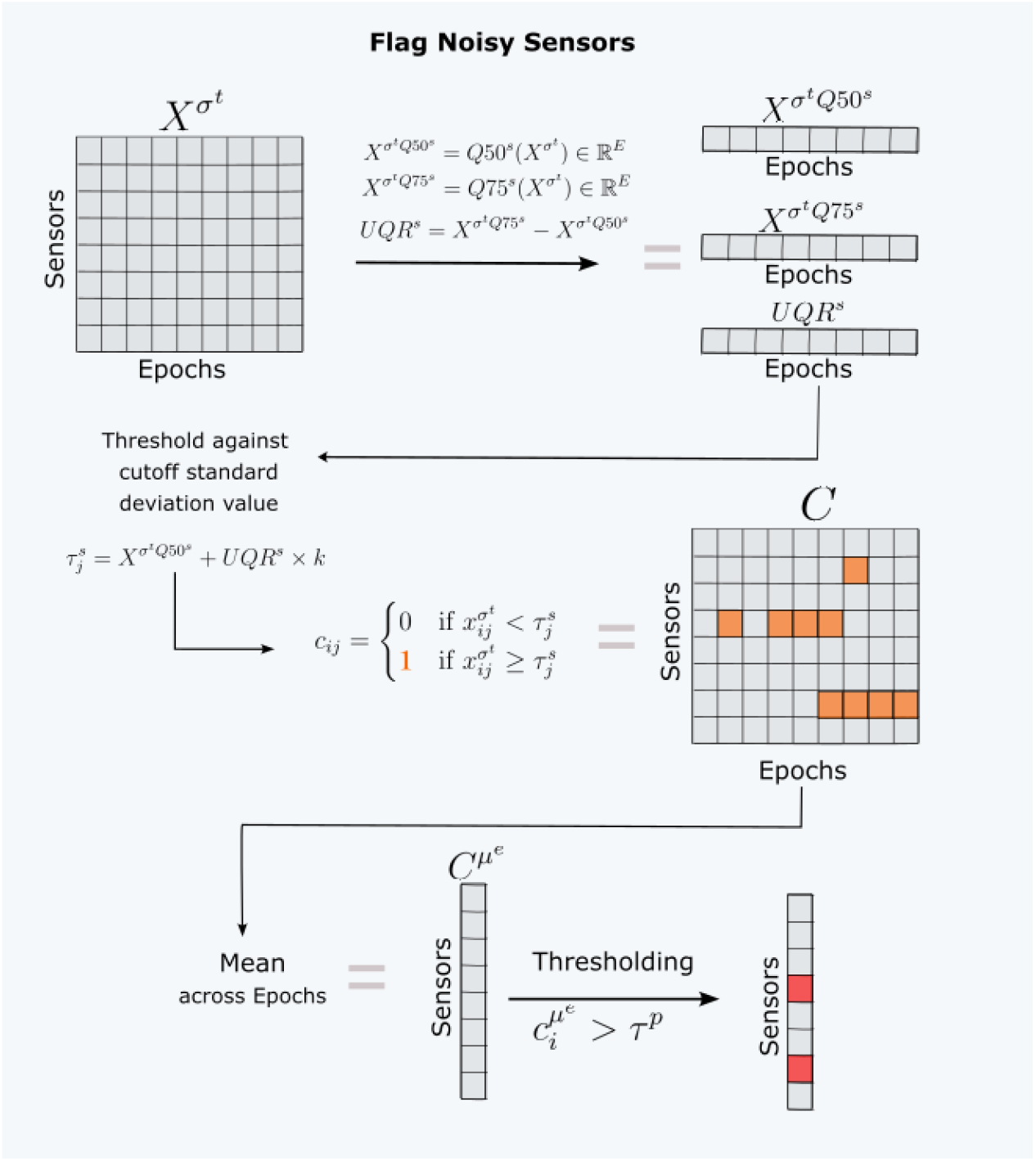
Schematic representation of the process used to flag noisy sensors.

### Flag Noisy Time-periods

The flag noisy time-periods step uses a similar approach as the flag noisy sensors step to identify and flag epochs with large, outlying voltage variance. This step is important because it can help to identify time periods that are contaminated by noise or large artifacts. For a detailed illustration of this step, see Supplementary Figure 2.

Again, the pipeline first takes the standard deviation of the input matrix *X* (which now excludes any sensors flagged in the previous step), resulting in a 2D matrix 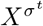. The pipeline then takes the 50th and 75th quantile of 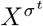 and calculates the upper quantile range across the epoch dimension, resulting in three vectors of size *S*_*𝒢*_ :

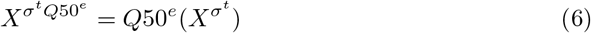

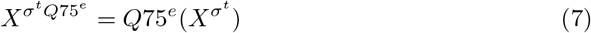

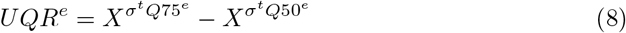

The sensor-specific thresholds for flagging epochs are then defined similarly as in (3), but swapping the role of the epoch and sensor dimensions:

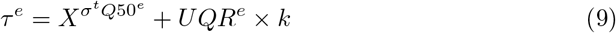

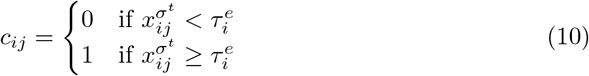

As before, each element of this matrix indicates whether a sensor *i* is an outlier at epoch *j*. In the previous step, the pipeline averaged across the epochs dimension to assess the percentage of epochs in which sensor *i* was an outlier. In this step, the pipeline averages across the sensor dimension of our indicator matrix *C* and obtains 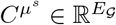, which is a vector of fractional numbers 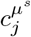, representing the percentage of sensors that are are outliers at each epoch *j*. Again, we use a consensus threshold *τ*^*p*^ as a cutoff point for determining if an epoch should be flagged. The epoch *j* is flagged as noisy if 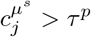. In other words, if more than *τ*^*p*^ sensors are outliers during epoch *j*, then epoch *j* is flagged as an artifact.

For any epoch that has been flagged by the pipeline, the start and end times are mapped back to the continuous EEG data and are saved within the MNE-Python Annotations structure to label corresponding time-periods.

### Filtering

By default, the pipeline applies a 1Hz highpass and a 100Hz lowpass to the data. The 1Hz low pass is recommended to avoid low frequency drifts from adversely affecting the independent component analyses (ICA) performed by the pipeline [13]. Further, the IClabel classifier used to categorize independent components was trained on data that had been bandpassed between 1Hz and 100Hz [8]. The users may alter these default settings in the configuration file. An additional notch filter can also be specified to remove line noise, which occurs at 60Hz or 50Hz depending on the region where the data was collected.

### Identify nearest neighbor sensors

This section does not describe a step but a preliminary computation required by a few subsequent steps that will be described afterward. The following steps in the PyLossless pipeline are designed to identify more fine-grained artifacts in the EEG. Due to the effect of volume conduction, neighboring EEG sensors are typically highly correlated with one another across time. The absence of correlation between neighboring sensors is indicative of non-neural artifacts, such as poor contact with the skin, or a defective sensor that produces a flat signal. Conversely, high correlations between neighboring sensors indicate bridging artifacts.

The previous steps for rejecting noisy sensors and epochs operated on a 2D matrix of dispersion values, specifically, the standard deviation across the samples dimension *t*. The following steps will operate on a matrix of correlation coefficients. In this section, we describe the procedure for defining this 2D matrix. Again, we start with the input matrix *X*, which now excludes previously flagged sensors and epochs. For each valid sensor *i* in *S*_*G*_, we select its *N* nearest neighbors, i.e. the *N* (default: N=3) sensors that are closest to it on the scalp. The specific number of neighboring sensors to include can be changed via the pipeline configuration file. We call the sensor *i* the origin and we use the symbol *ŝ*_*l*_ (with *l* ∈ {1, 2, …, *N*}) to refer to its nearest neighbors.

For each epoch *j*, we calculate the correlation coefficient 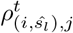 between origin sensors *i* and their neighbors *ŝ*_*l*_ across the samples dimension *t*, returning a 3D matrix of correlation coefficients:

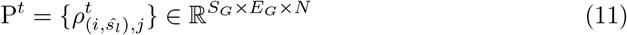

Then, we identify the sensor with the maximal correlation across the “neighbor” dimension *n* (see Fig 4) as:

**Fig 4.**
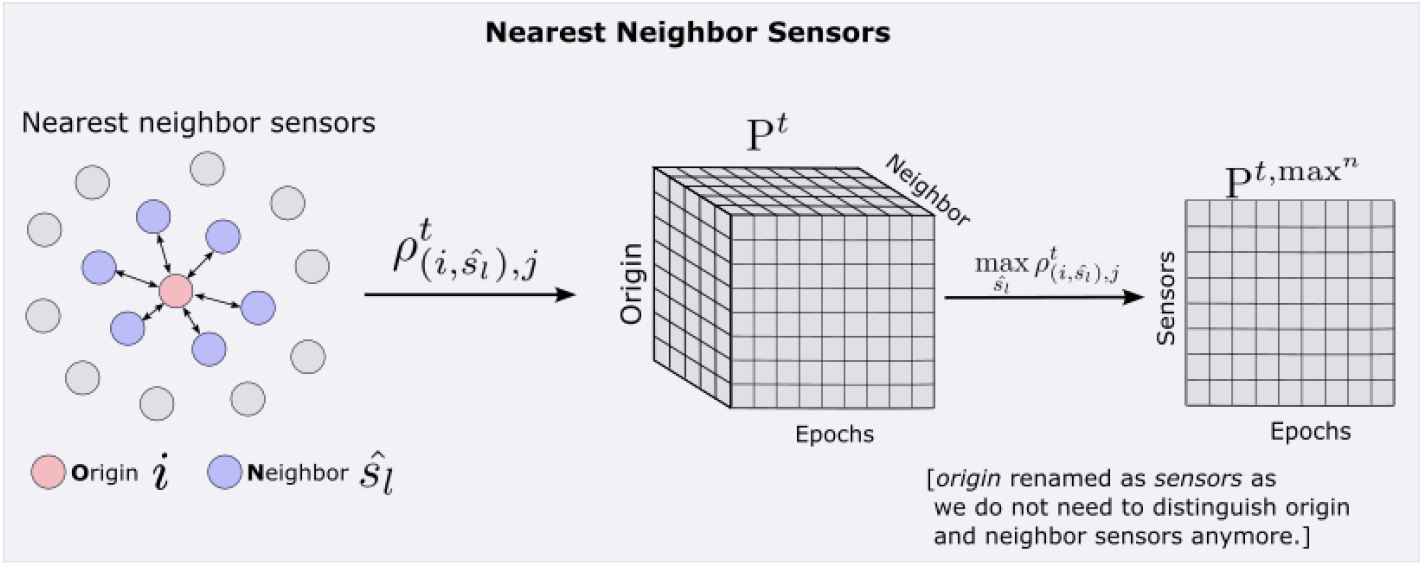
Visual representation of the procedure for identifying nearest neighbor sensors. See the text for the definition of the mathematical symbols.

**Fig 5.**
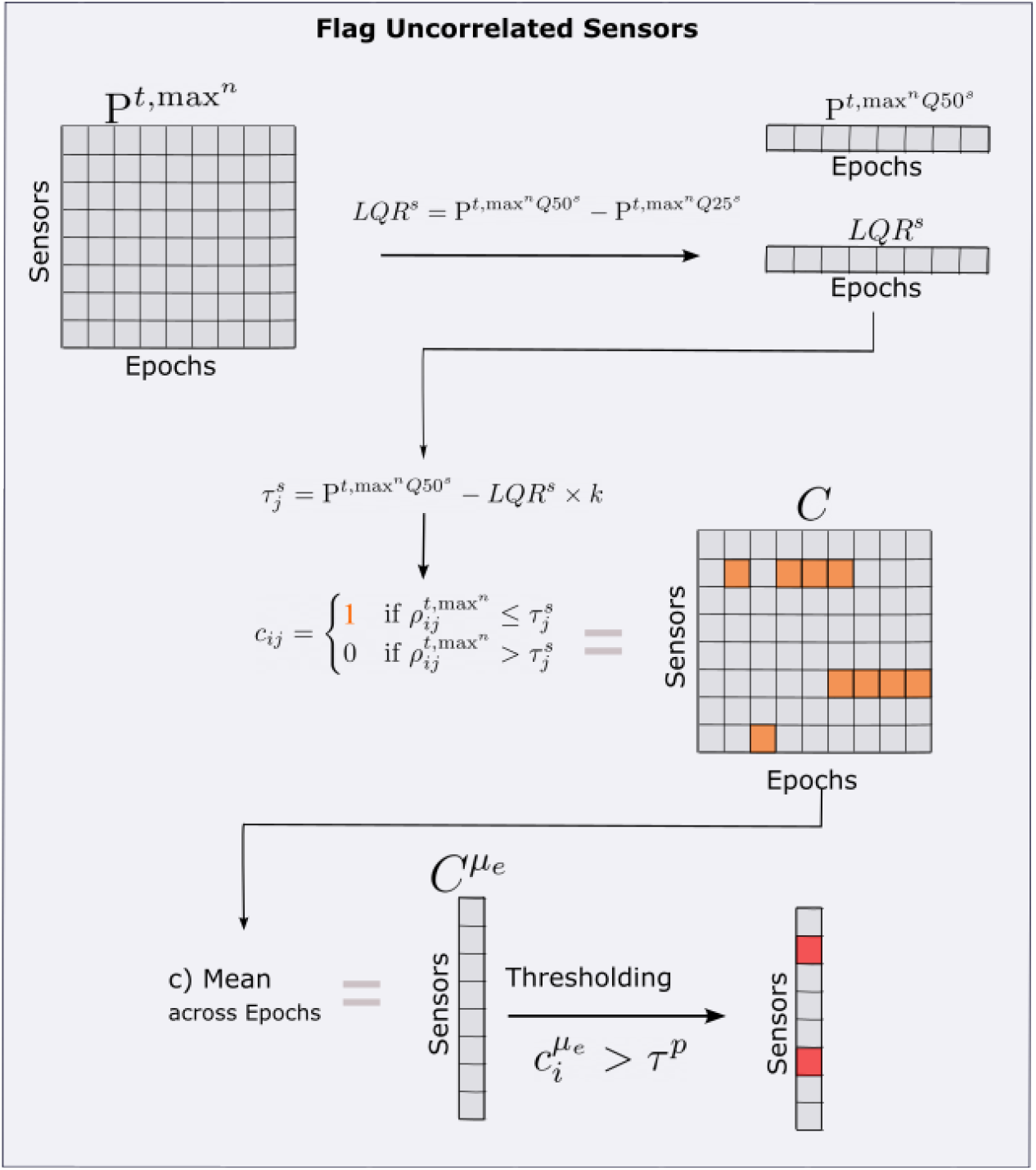
Visual representation of the flag uncorrelated sensors step. See the text for the definition of the mathematical symbols.

**Fig 6.**
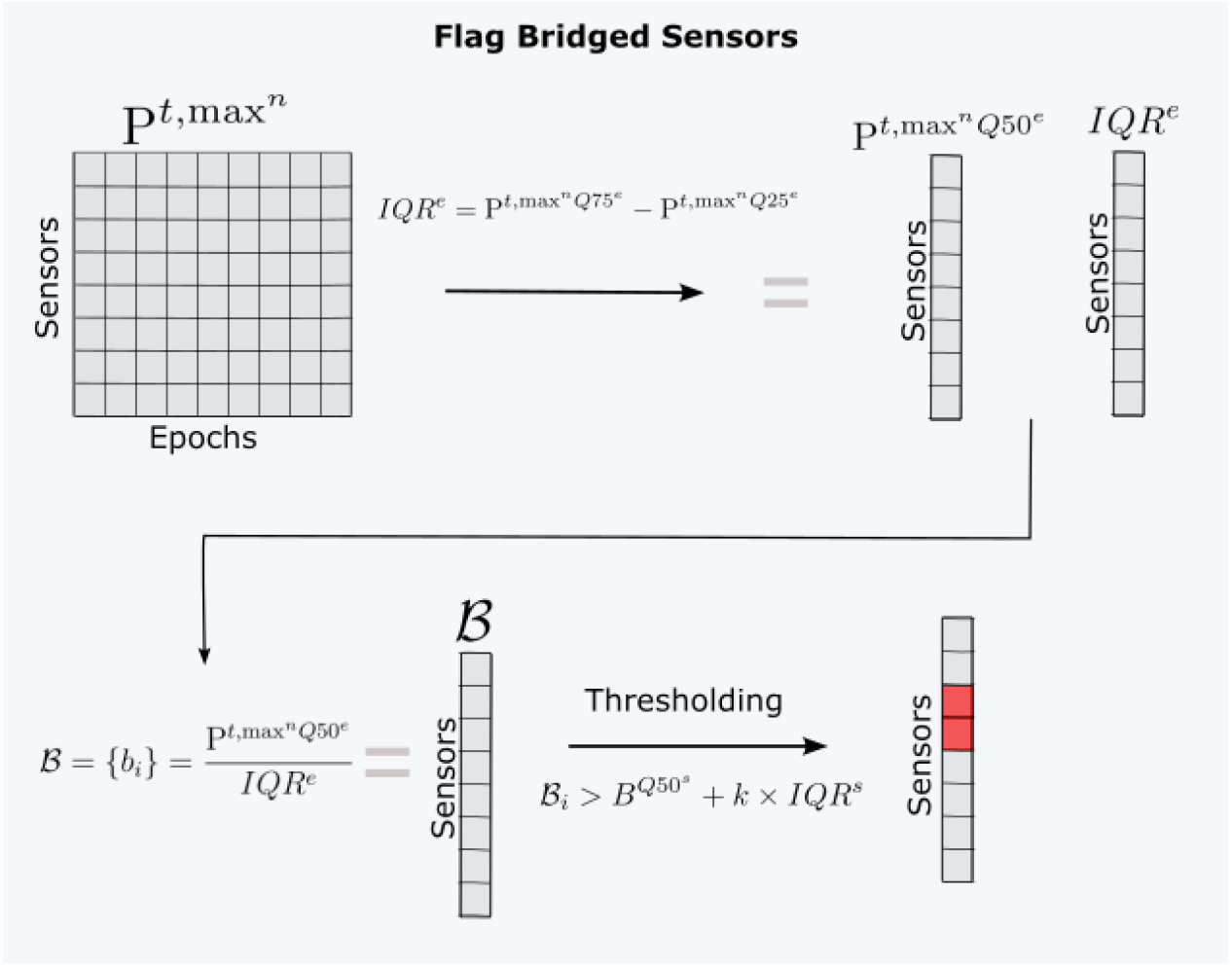
Schematic representation of the process used to flag bridged sensors.

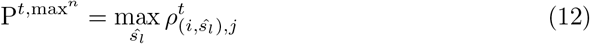

The next few steps use this 2D matrix is used as an input.

### Flag uncorrelated sensors

This step is designed to identify sensors that have an unusually low correlation with neighboring sensors. Due to the effect of volume conduction, a single source of neural activity is often picked up by several sensors on the scalp. Thus, the signals of neighboring EEG sensors are typically highly correlated across time. When the signal of an EEG sensor is uncorrelated with the signals of neighboring sensors, it is indicative of an artifact caused by poor electrode-skin contact [1]. The operations involved in this step are similar to those of the *flag noisy sensors* step, except we use maximal nearest neighbor correlations instead of dispersion and the left instead of the right tail of the distribution to set the threshold for outliers. First, we take the 25th and 50th quantiles of 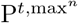, across the sensors dimension, and calculate the lower quantile range *LQR*^*s*^. This results in vectors 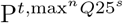 and 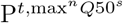, each of size *E*_*𝒢*_. The lower quantile range (*LQR*) is defined as:

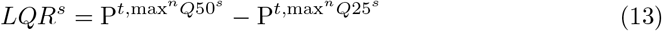

Next, we define epoch-specific thresholds for rejecting sensors

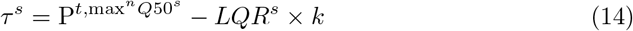

with 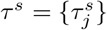, where 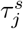 denote a threshold specific to epoch *j*.

Then, the pipeline compares each column *j* of the 2D matrix of maximal correlation coefficients 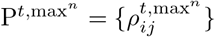 to the threshold 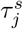 resulting in the indicator matrix *C* ∈{0, 1}^*S×E*^ ={*c*_*ij*_}. If the correlation coefficient of sensor *i* at epoch *j* is below the epoch-specific threshold 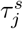, then the epoch *j* is considered as containing an artifact for channel *i* and is labeled as such in the indicator matrix:

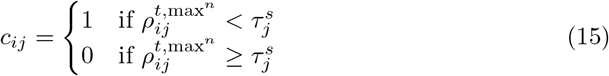

Similar to the flag noisy sensors step, the pipeline averages across the epochs dimension of our indicator matrix *C* and obtains 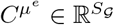, representing the percentage of epochs in which sensor *i* is uncorrelated with its neighbors. We define a consensus threshold *τ*^*p*^ as a cutoff point for determining if a sensor should be flagged. The sensor *i* is flagged if 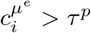.

### Flag Bridged Sensors

This section describes the operations performed for detecting and flagging bridged sensors. Bridging of sensors happens when an abnormally high conductivity is introduced between two sensors, e.g., if electrolyte paste “bridges” two electrodes. First, the pipeline takes the 25th, 50th, and 75th quantiles across the epochs dimension of input matrix 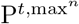, and calculates the inter quantile range (IQR):

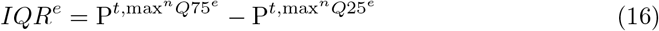

For each sensor *i*, we divide the median across epochs by the *IQR* across epochs. Bridged sensors should have a high median correlation but a low *IQR* of the correlation, i.e., a bridged channel has a consistently high temporal correlation with the channel it is bridged with. We call this measure the bridge indicator, which is calculated as follows:

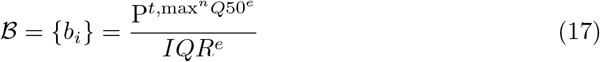

Next, we define a bridging threshold. We take the 25th, 50th, and 75th quantiles of ℬ and calculate the scalar-valued *IQR* across sensors (*IQR*^*s*^). A sensor *i* is bridged if 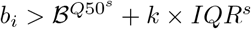. Any sensor selected by this threshold is flagged as “bridged”.

Finally, a robust average reference is again applied to the data to exclude any sensor that has been flagged as noisy, uncorrelated, or bridged.

### Flag the rank sensor

Because the pipeline uses an average reference before the ICA decomposition, it is necessary to account for rank deficiency (i.e., every sensor in the montage is linearly dependent on the other channels due to the common average reference). To account for this, PyLossless flags the sensor (out of the remaining good sensors) with the highest median maximal correlation coefficient with its neighbors (across epochs):

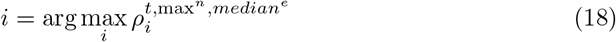

This sensor has the least unique time-series out of the remaining set of good sensors *S*_*G*_ and is flagged by the pipeline as “rank”. Note that this sensor is not flagged because it contains artifact, but only because one of the remaining sensors needs to be removed to address rank deficiency before ICA decomposition is performed. By choosing this sensor, we are likely to lose little information because of its high correlation with its neighbors. This sensor can be reintroduced by interpolation after the ICA has been applied for artifact corrections.

### Flag uncorrelated epochs

This step is designed to identify periods in which many sensors are uncorrelated with neighboring sensors. Because EEG signals of the neural sources are heavily smeared as their electrical potential crosses the relatively resistive skull, scalp EEG has a low spatial frequency and, hence, exhibits a relatively high correlation between neighboring sensors. Neighbor electrodes capturing uncorrelated signals are generally due to artifacts, e.g., a very high impedance of one of the two sensors causing it to capture mostly ambient electromagnetic noise. Again, we calculate the 25th and 50th quantile of 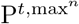, across the epochs dimension, and calculate the lower quantile range *LQR*^*s*^. This results in vectors 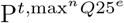 and 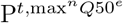 of size *S*_*𝒢*_. As for previous steps, we define sensor-specific thresholds for flagging epochs:

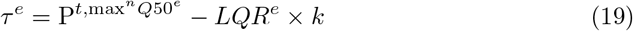

And the corresponding indicator matrix:

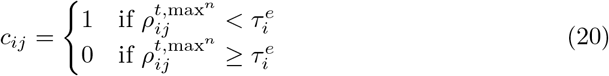

We average the indicator matrix across sensors and obtain a vector 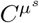 that we use to flag uncorrelated epochs using the following criterion: 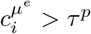.

### Initial ICA and flagging of outlying independent component (IC)

To remove artifact components from the EEG, PyLossless runs two subsequent ICA decompositions. The first decomposition is a preprocessing step meant to remove non-stationary segments of data so that the second ICA can classify more accurately ICs according to their likely origin (e.g., brain, cardiac, ocular, etc.). PyLossless uses the built-in ICA routines available in MNE-Python. Because the initial ICA decomposition is only used to identify periods with clear non-stationary activity and does not require very accurate decomposition, the pipeline uses by default the fastICA algorithm [14]. This setting can be changed via the pipeline configuration file.

The IClabel classifier (and ICA decompositions in general) performs better in the absence of non-stationarity in the data [13]. To detect and remove such non-stationary periods, PyLossless repeats the operations described in the flag noisy epochs step, but replacing the *X* matrix by a similar 3D matrix 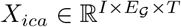 using IC activation time-courses rather than scalp EEG data and where *I* is the set of independent components. In short, using the terminology defined in the flag noisy epochs step, the pipeline flags as flag noisy ICs any epoch *j* where more than *τ*^*p*^ percent of ICs contained outlying dispersion values.

### Final ICA and IC classification

For the final ICA decomposition, the pipeline uses the Extended Infomax algorithm because it generally produces reliable decompositions, and the IClabel classifier was trained on ICA produced with this algorithm [15]. The resulting ICA decomposition is processed by the IClabel classifier [16, 17], implemented via the python package MNE-ICALabel [8], which uses a neural network to categorize each independent component as either neural, muscle artifact, eye blink, heartbeat, line noise, channel noise, or other. Additionally, a confidence rating is provided for each component label.

### Saving the pipeline output

After the automated pipeline has terminated, the annotated EEG, ICA data, and pipeline flags are saved in a *derivatives* subdirectory [18] (i.e., project root/derivatives/pylossless, where project root is the original BIDS-compliant directory that was passed into the pipeline) using the format and file structure defined by the BIDS-EEG specification [5]. The EEG data are saved in European Data Format (EDF; [19]), as supported by BIDS. The ICA data are saved using MNE-Python’s built-in ICA saving functionality. The pipeline decisions on sensors are saved as tab-separated text files (TSV), and the MNE-ICALabel decisions regarding independent component categorization are saved in a comma-separated text file (CSV). The pipeline decisions about flagged time periods are also saved as annotations directly within the EEG data. Finally, the PyLossless pipeline saves a YAML configuration file named “rejection policy.yaml”. This file specifies how to use the various pipeline flags in the destructive phase of the data preprocessing and needs to be provided to apply analysis-specific cleaning procedures. Copies of this file can be made, and their rules modified to adjust in a traceable way how the sharable data state (as defined in Fig. 1) is transformed into a preprocessed dataset, dependent on the specific requirements of the different analyses.

### Quality Control Review

After the automated step of the pipeline is completed, the users can review the PyLossess decisions via a streamlined QCR dashboard. This dashboard presents the user with information about the EEG data, ICA, and the decisions made by the pipeline. It is served using Plotly-Dash, an open-source Python package that uses JavaScript bindings to create interactive graphics. This dashboard can be opened in the user’s default browser (Fig 7) from a terminal using the command pylossless qc.

**Fig 7.**
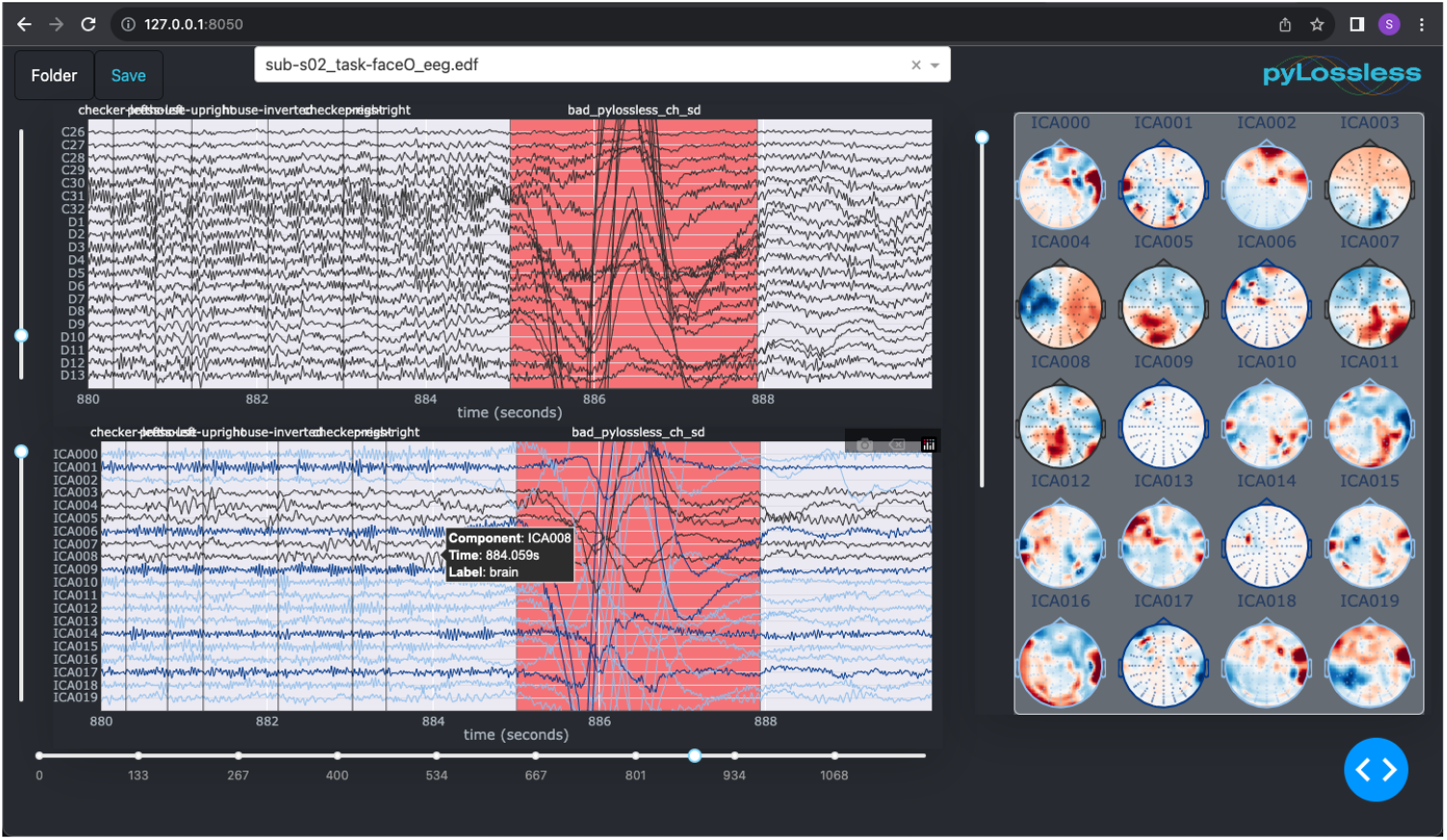
The PyLossless QCR Dashboard. The top-left panel displays the continuous EEG data, and the bottom-left panel displays the IC activations. The right panel displays the topomaps for each IC.

The QCR dashboard allows users to select PyLossless output files via a directory selection dialog and a file selection dropdown menu. Once a file is selected, the continuous EEG (Fig 7, top-left window), the ICs from the final ICA (Fig 7, bottom-left window), and the topographic plots corresponding to the ICs (Fig 7, right window) are displayed.

The user can assess and modify the decisions of the pipeline regarding the flagging of channels, artifacts, and IC, as well as the overall quality of the final ICA decomposition. When hovering over an IC activation or a topographic plot, a hover label displays the corresponding IClabel classification and the associated confidence rating. Further, the IC activations are color-coded by category (e.g., brain, muscle artifact, etc.). Sensors and ICs flagged by the pipeline are grayed out. Time periods flagged by the pipeline are covered by red rectangles, with text over them designating the annotation label (e.g., *BAD PyLL noisy*).

The user can confirm the decisions of the automated pipeline by not altering the flags, or they can revise, remove, or add new flags. Users can flag additional sensors or ICs by clicking on them. Any sensor or IC flagged by the user is given the label BAD PyLL manual”. Likewise, users can click on flagged sensors or independent components to override their flag status and consider flagged data valid. Users can add new flagged time periods by clicking and dragging over a sensor or independent time-series plot. This click-drag event results in a new red rectangle over the designated time period, with the annotation label *BAD PyLL manual*. Finally, the user can revise or remove any time-based annotations via the user interface.

Once the user has finished the QCR, they can save the updated data to the disk. Since the QCR process does not change the EEG signals, but only the pipeline flags and the annotations associated with the data, the original files are overwritten when saving the QCR output. Overwriting instead of duplicating ensures that the pipeline output does not take too much disk storage, as continuous EEG files are often hundreds of megabytes, or even gigabytes, in size.

### Applying the PyLossless decisions

By design, PyLossless does not alter the input EEG data. To apply the pipeline’s decisions, PyLossless provides a `RejectionPolicy` class, which specifies the flags and IC categories that should be removed from the EEG data. The rejection policy can be loaded using the provided `read rejection policy` function, which loads the rejection policy YAML configuration file from the disk. By default, the PyLossless rejection policy will add any flagged sensors as bad sensors to the MNE-Python info structure of the continuous raw EEG. For ICs, the pipeline will subtract from the EEG any IC labeled as a category other than “brain” or “other” with a confidence rating of at least 30%. This rejection policy is fully configurable by the user, for example allowing not to remove EOG components for a study aiming to use these components in its analysis.

## Results

We confirmed that EEG recordings cleaned by PyLossless are of equal or better quality than those from the MATLAB EEG-IP-Lossless pipeline by comparing the performance of these two pipelines using the same open-access dataset previously used to assess the performance of EEG-IP-Lossless [4]. This dataset and the corresponding experimental task are described in [20]. In short, a visual processing task using inverted faces and inverted houses as stimuli was used with 10 participants. The EEG was epoched to the stimuli onsets, using the preceding 200 ms as a baseline, and the following 600 ms as the response to the stimulus. One of the main findings of the publication was related to an ERP difference in inverted face stimuli as compared to inverted house stimuli. The authors provided a graphic displaying the time-courses of the average global field amplitude (GFA) for the inverted face, the inverted house, and the difference between these two conditions. The between-condition difference was computed by resampling the epochs from each condition and computing the corresponding curves 1000 times (Figure 8, grey lines). As can be seen, the GFA waves from the data cleaned by PyLossless (Figure 7, bottom panel) are nearly identical to the original graphic from the publication (Figure 8, top panel), suggesting that PyLossless clean EEG as well if not better than the previous MATLAB version (as shown by larger ERP amplitudes), while benefiting from a significant improvement in usability.

**Fig 8.**
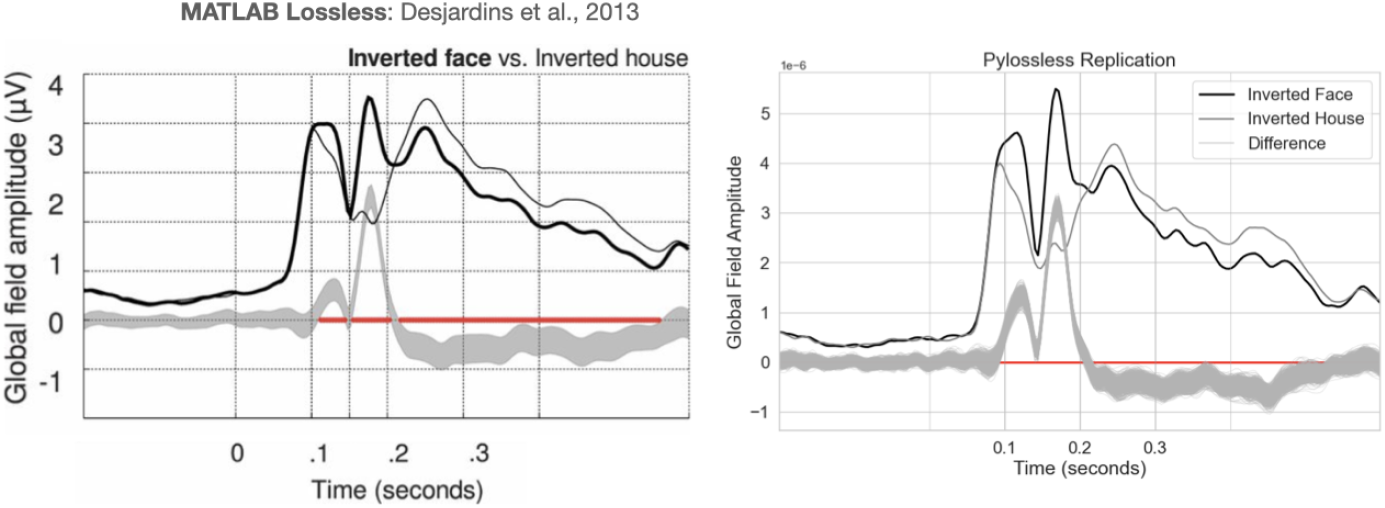
Replication of the Global Field Amplitude figure from Desjardins and colleagues (2013) [20].

**Fig 9.**
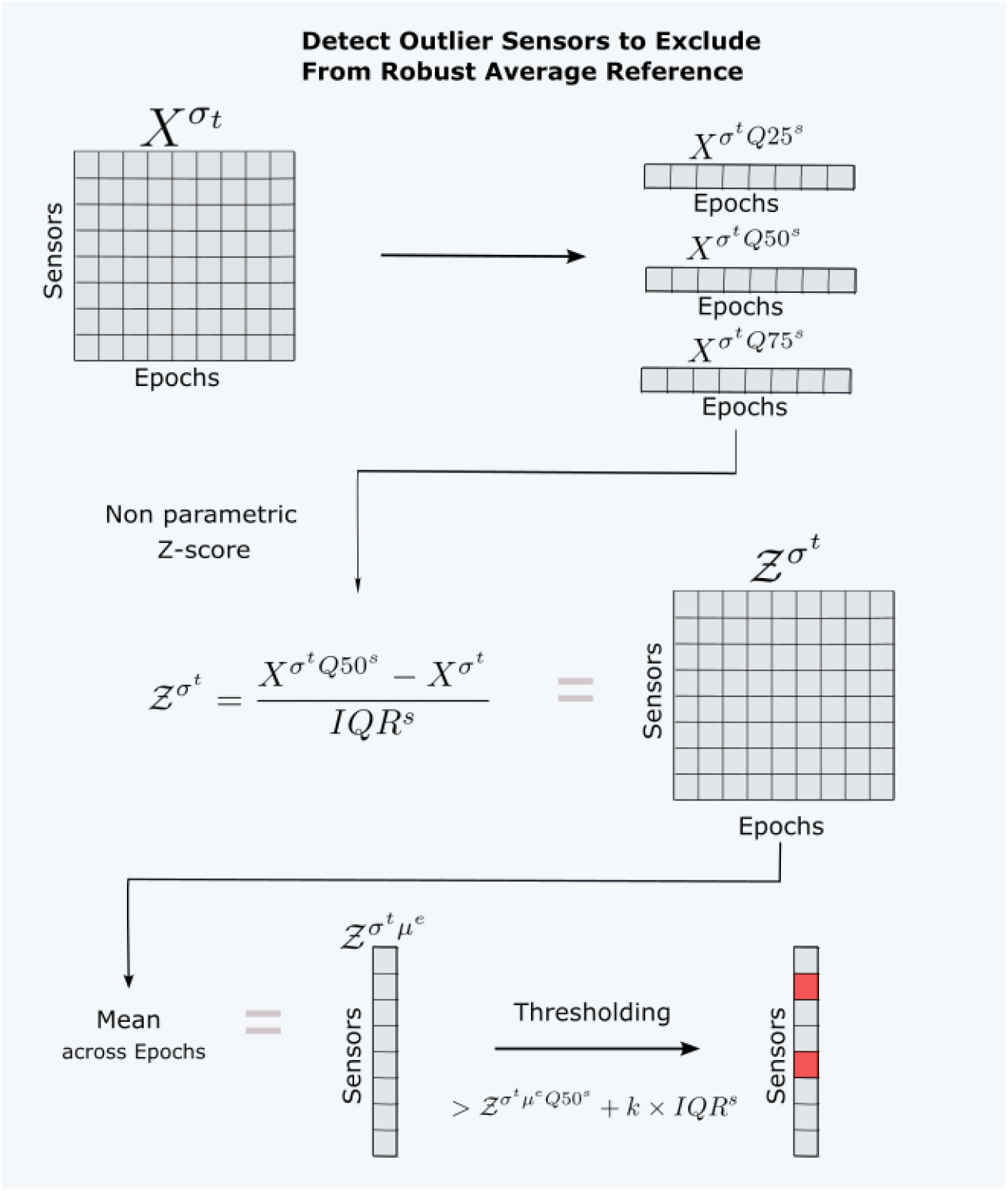
Schematic representation of how PyLossless detects sensors to exclude from the average reference.

## Discussion

Open-source, large EEG datasets are becoming increasingly common [21], and there is a need for procedures that allow researchers to readily access cleaned EEG data, with flexibility regarding artifact rejection. The aim of this project was to provide robust open-source software for cleaning EEG that provides a sharable annotated data state that preserves the original data and can be used in a traceable way to produce alternate preprocessing at the time of analysis. For example, researchers may wish to apply or disregard ICA, or to remove only specific eye-blink components. They may want to epoch EEG into 1- or 10-second segments, depending on the analysis they will carry out (e.g., for connectivity analyses in resting-state recordings, the optimal epoch length depends on the different functional connectivity measures). In an ideal scenario, researchers would not need to individually process the data due to slight differences in their desired output. Pylossless aims to make this possible, with the goal of making it easier for research teams or communities to collaborate on shared EEG datasets.

To achieve this objective, it was important to retain the original continuous state, as opposed to managing artifact rejections only on epoched data. Building this pipeline on top of MNE-Python provides a key advantage, as MNE natively supports annotating the continuous EEG structure. Further, Python is now the most widely used programming language and has a mature scientific library stack and a growing community [22]. MNE-Python is also a very healthy package (i.e., a 93/100 Package Health Score according to the Snyk Abvisor [23]), ensuring sustainable development. Thus, we believe that providing this pipeline in Python will increase its accessibility to the growing Python scientific community.

### Encouraging Data Review

PyLossless provides a QCR dashboard to help the users review the pipeline’s decisions. The purpose of the QCR dashboard is to encourage users to engage with the data during the initial preprocessing stage. After running the automated pipeline, users can review the pipeline output on individual recordings to confirm that noisy sensors and time periods have been flagged and that the ICA decomposition sufficiently isolated spatially stationary artifacts (EOG, ECG, etc.) from neural signals. Ultimately, the QCR dashboard aims to support expert confirmation of Pylossless preprocessing quality before the datasets are used (or reused by other researchers) for hypothesis testing.

### Improvements in software design and stability

PyLossless is the natural evolution of the MATLAB EEG-IP-Lossless pipeline [4]. We significantly improved on this prior work by further developing the ideas pushed forward by [4] and repackaging the EEG-IP-Lossless functionalities into robust and accessible software. In doing so, we clarified foundational concepts, made PyLossless simple to install and use, and provided thorough documentation, including the API definition and tutorials. We adhered to programming best practices, including thorough testing and continuous integration (CI), as implemented within the open-source Python ecosystem. Finally, unlike its MATLAB predecessor, PyLossless can be used easily on remote compute clusters or services like Google Colab because it does not require proprietary software like MATLAB and its graphical component is web-based. This reliance on web technologies allows researchers to use the pipeline on remote compute clusters or Platform As A Service (PAAS) providers such as Google Cloud Engine or Microsoft Azure, unlocking additional computational power for processing in parallel large datasets.

In particular, we believe that the PyLossless QCR dashboard is a significant contribution to the EEG research community, with the Dash-Plotly-based QCR dashboard allowing users to visualize preprocessed EEG on almost any platform. Further, because Plotly-Dash is designed to share graphics via the web, the opportunity is available for researchers to share visualizations of EEG from open datasets via the web. Future developments of PyLossless may see the QCR component migrated to a stand-alone software package, to further develop features for researchers to visualize their data in web frameworks, and collaboratively interact with data. We are also considering implementing more visual diagnostic functions to help researchers efficiently assess the quality of their data and the performance of preprocessing procedures. We would also like to port the EEG-IP-Lossless average referencing approach that uses a common interpolated EEG grid on a standard head surface to increase the comparability of results obtained on datasets using nets with significantly different head coverage.

## 1 Acknowledgments

The authors acknowledge the following sources of Funding: Christian O’Reilly is supported through startup funds from the University of South Carolina. Mayada Elsabbagh is partially supported through the Fonds de Recherche du Québec (FRQS; grant number 34734).

## Supporting information

### Robust Average Reference

The robust average reference is performed before the first pipeline step and between most pipeline steps. This routine uses a standard deviation-based metric to identify sensors with outlying voltage variance across time and leaves them out of the reference operation. For this routine, a 3D matrix standard deviation is taken across dimension *t* of input matrix *X*, resulting in 2D matrix 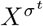. Then, the pipeline computes 25th, 50th, and 75th quantiles across the sensors dimension of 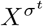, and the Inter-Quantile-Range (IQR) is calculated:

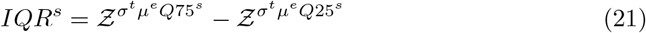

Then, a non-parametric Z-score is computed from these vectors:

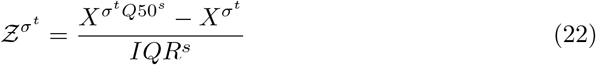

Each sensor *i* is left out of the rereferencing procedure if it has previously been flagged or if 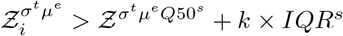 where 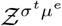 is the mean of 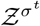 across epochs. This process is illustrated in Supplementary Figure 9.

### Flag Noisy Time Periods

The *flag noisy time periods* step is very similar to the *flag noisy sensors* step previously described, but exchanging the dimensions along which the various operations are performed. This process is described in Supplementary Figure 10.

**Fig 10.**
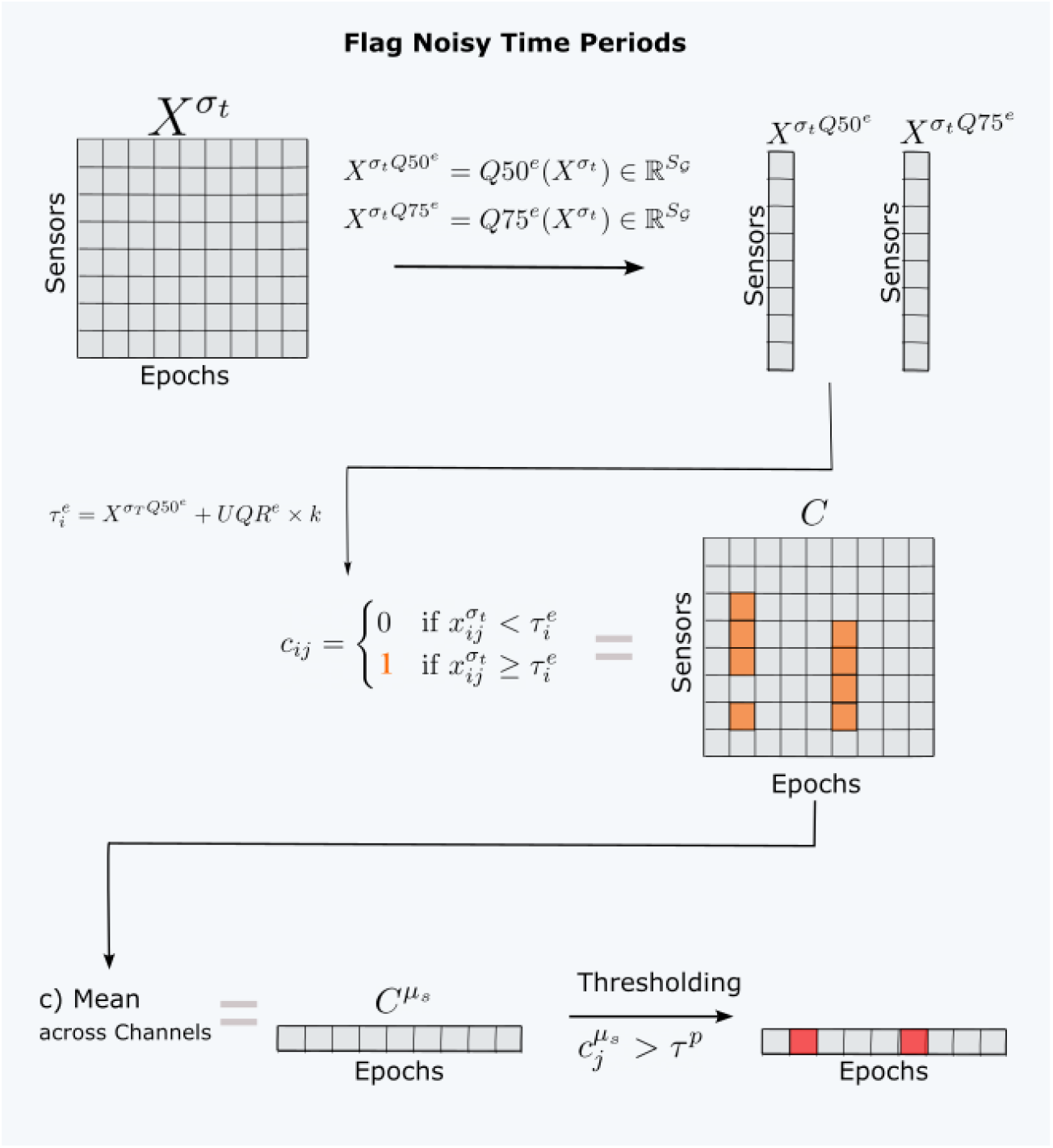
Schematic representation of the process used to flag noisy time periods.

## References

1. Bigdely-Shamlo N, Mullen T, Kothe C, Su KM, Robbins KA. The PREP pipeline: standardized preprocessing for large-scale EEG analysis. Frontiers in Neuroinformatics. 2015;9. doi:10.3389/fninf.2015.00016.

2. Gabard-Durnam LJ, Mendez Leal AS, Wilkinson CL, Levin AR. The Harvard Automated Processing Pipeline for Electroencephalography (HAPPE): Standardized Processing Software for Developmental and High-Artifact Data. Frontiers in Neuroscience. 2018;12. doi:10.3389/fnins.2018.00097.

3. Jas M, Engemann D, Raimondo F, Bekhti Y, Gramfort A. Automated rejection and repair of bad trials in MEG/EEG. In: 2016 International Workshop on Pattern Recognition in Neuroimaging (PRNI). Trento, Italy: IEEE; 2016. p. 1–4. Available from: http://ieeexplore.ieee.org/document/7552336/.

4. Desjardins JA, van Noordt S, Huberty S, Segalowitz SJ, Elsabbagh M. EEG Integrated Platform Lossless (EEG-IP-L) pre-processing pipeline for objective signal quality assessment incorporating data annotation and blind source separation. Journal of Neuroscience Methods. 2021;347:108961. doi:10.1016/j.jneumeth.2020.108961.

5. Pernet CR, Appelhoff S, Gorgolewski KJ, Flandin G, Phillips C, Delorme A, et al. EEG-BIDS, an extension to the brain imaging data structure for electroencephalography. Scientific Data. 2019;6(1):103. doi:10.1038/s41597-019-0104-8.

6. Gramfort A. MEG and EEG data analysis with MNE-Python. Frontiers in Neuroscience. 2013;7. doi:10.3389/fnins.2013.00267.

7. Appelhoff S, Sanderson M, Brooks TL, van Vliet M, Quentin R, Holdgraf C, et al. MNE-BIDS: Organizing electrophysiological data into the BIDS format and facilitating their analysis. Journal of Open Source Software. 2019;4(44):1896. doi:10.21105/joss.01896.

8. Li A, Feitelberg J, Saini AP, Höchenberger R, Scheltienne M. MNE-ICALabel: Automatically annotating ICA components with ICLabel in Python. Journal of Open Source Software. 2022;7(76):4484. doi:10.21105/joss.04484.

9. Harris CR, Millman KJ, van der Walt SJ, Gommers R, Virtanen P, Cournapeau D, et al. Array programming with NumPy. Nature. 2020;585:357–362. doi:10.1038/s41586-020-2649-2.

10. Virtanen P, Gommers R, Oliphant TE, Haberland M, Reddy T, Cournapeau D, et al. SciPy 1.0: fundamental algorithms for scientific computing in Python. Nature Methods. 2020;17(3):261–272. doi:10.1038/s41592-019-0686-2.

11. The pandas development team. pandas-dev/pandas: Pandas;. Available from: https://github.com/pandas-dev/pandas.

12. PyLossless contributors. PyLossless Documentation; 2023. Available from: https://pylossless.readthedocs.io.

13. Winkler I, Debener S, Muller KR, Tangermann M. On the influence of high-pass filtering on ICA-based artifact reduction in EEG-ERP. In: 2015 37th Annual International Conference of the IEEE Engineering in Medicine and Biology Society (EMBC). Milan: IEEE; 2015. p. 4101–4105. Available from: http://ieeexplore.ieee.org/document/7319296/.

14. Ablin P, Cardoso JF, Gramfort A. Faster Independent Component Analysis by Preconditioning With Hessian Approximations. IEEE Transactions on Signal Processing. 2018;66(15):4040–4049. doi:10.1109/TSP.2018.2844203.

15. Lee TW, Girolami M, Sejnowski TJ. Independent Component Analysis Using an Extended Infomax Algorithm for Mixed Subgaussian and Supergaussian Sources. Neural Computation. 1999;11(2):417–441. doi:10.1162/089976699300016719.

16. Pion-Tonachini L, Kreutz-Delgado K, Makeig S. ICLabel: An automated electroencephalographic independent component classifier, dataset, and website. NeuroImage. 2019;198:181–197. doi:10.1016/j.neuroimage.2019.05.026.

17. Pontifex MB, Gwizdala KL, Parks AC, Billinger M, Brunner C. Variability of ICA decomposition may impact EEG signals when used to remove eyeblink artifacts: ICA variability. Psychophysiology. 2017;54(3):386–398. doi:10.1111/psyp.12804.

18. BIDS Contributors. Storage of derived datasets;. Available from: https://bids-specification.readthedocs.io/en/stable/02-common-principles.html#storage-of-derived-datasets.

19. Kemp B, Värri A, Rosa AC, Nielsen KD, Gade J. A simple format for exchange of digitized polygraphic recordings. Electroencephalography and Clinical Neurophysiology. 1992;82(5):391–393. doi:10.1016/0013-4694(92)90009-7.

20. Desjardins JA, Segalowitz SJ. Deconstructing the early visual electrocortical responses to face and house stimuli. Journal of Vision. 2013;13(5):22–22. doi:10.1167/13.5.22.

21. Markiewicz CJ, Gorgolewski KJ, Feingold F, Blair R, Halchenko YO, Miller E, et al. The OpenNeuro resource for sharing of neuroscience data. eLife. 2021;10:e71774. doi:10.7554/eLife.71774.

22. TIOBE. TIOBE Index, July 2023; 2023. https://www.tiobe.com/tiobe-index/.

23. Snyk Advisor. Snyk Advisor: mne package health score; 2023. Available from: https://snyk.io/advisor/python/mne.

